# CXCL12, SCF, and eotaxin are prognostic serum biomarkers in gastric cancer

**DOI:** 10.1101/2025.08.17.670705

**Authors:** Jefim Brodkin, Tuomas Kaprio, Harri Mustonen, Alli Leppä, Arto Kokkola, Marko Salmi, Sirpa Jalkanen, Caj Haglund, Camilla Böckelman

**Affiliations:** Translational Cancer Medicine Research Program, Faculty of Medicine, University of Helsinki, Helsinki, Finland; Department of Surgery, University of Helsinki and Helsinki University Hospital, Helsinki, Finland; MediCity Research Laboratory and Institute of Biomedicine, University of Turku, Turku, Finland; InFLAMES Flagship, University of Turku, Turku, Finland; Department of Pathology, University of Helsinki and Helsinki University Hospital, Helsinki, Finland

**Author notes:** Correspondence to: Jefim Brodkin, MD, doctoral researcher, Translational Cancer Medicine Research Program, Faculty of Medicine, University of Helsinki, Haartmaninkatu 4, PO Box 340, FIN-00029 HUS Finland. Shared last authorship (Caj Haglund, Camilla Böckelman).

**Keywords:** Gastric cancer, Survival, CXCL12, SCF, Eotaxin

## Abstract

Gastric cancer is the fifth most common cancer worldwide and the fifth leading cause of cancer-related death. Its poor prognosis is primarily due to a late diagnosis and a lack of effective treatments for advanced disease. We aimed to identify new prognostic serum biomarkers to aid clinical decision-making.

Our patient cohort consisted of 240 individuals who underwent surgery for histologically verified gastric adenocarcinoma in the Department of Surgery, Helsinki University Hospital, between 2000 and 2009. To determine serum protein concentrations of cytokines and growth factors, we utilized Bio-Rad’s premixed Bio-Plex Pro Human Cytokine 27-plex and 21-plex assay kits.

Among the 48 biomarkers analyzed, three emerged as statistically significant prognostic markers for disease-specific survival using the Cox proportional hazards univariate analysis: C-X-C motif chemokine ligand 12 (CXCL12) (hazard ratio [HR] 0.39, 95% confidence interval [CI] 0.23–0.63, *p*<0.001), stem cell factor (HR 0.38, 95%CI 0.19–0.77, *p*=0.007), and eotaxin (HR 0.57, 95%CI 0.37–0.89, *p*=0.013).

Multivariate survival analysis revealed that, among the 48 biomarkers analyzed, CXCL12 and eotaxin served as independent prognostic markers among gastric cancer patients. The prognostic effect of inflammatory serum biomarkers in gastric cancer could provide new insights into the immunological microenvironment of disease.

## Introduction

Gastric cancer (GC) is a common cancer worldwide. Although its incidence has fallen in the Western world, GC remains one of the leading causes of cancer-related deaths, with the fifth highest incidence and fifth most common cause for cancer-related death globally [1].

The incidence of GC has decreased in Finland and other Western countries in recent decades. However, the prognosis of GC remains quite poor: the five-year survival rate in Finland from 2020 to 2022 was just above 30% [2], similar to other Western countries that do not screen for GC. Its poor prognosis is primarily due to a late diagnosis and ineffective treatments for metastasized disease. To improve overall survival, new treatments and earlier diagnostics are needed.

Patient age and stage are known prognostic markers for GC. Additionally, patients with a diffuse histology according to the Laurén classification [3] experience a worse prognosis. Novel molecular subtypes such as those introduced by The Cancer Genome Atlas (TCGA) [4] and the Asian Cancer Research Group (ACRG) [5] may serve as potential prognostic markers.

Various serum biomarkers, such as C-reactive protein (CRP), carcinoembryonic antigen (CEA), and carbohydrate antigen 19-9 (CA19-9), are used in the diagnosis and follow-up of GC patients. However, their prognostic value remains unclear. For instance, Lu et al. [6] showed that patients with high pre-operative and/or post-operative CRP levels exhibit a worse prognosis. However, some other studies have shown no or a weak effect on survival [7, 8]. A meta-analysis found an elevated CRP level in patients with a worse prognosis [9]. However, Fent et al. [10] demonstrated that patients with high CEA levels have a worse survival, but that CA19-9 was not a prognostic factor. Levels of CEA and CA19-9 can be used to assess the effect of neoadjuvant treatment, whereby normalization of their levels is indicative of a better survival [11]. Additionally, GC patients with recurring disease and a worse prognosis had higher CEA levels, whereas no difference in the CA19-9 levels was observed [12].

Cancer and inflammation are intertwined, such that GC is an example of an infection-driven cancer where most cases are associated with *H. pylori* or Epstein–Barr virus (EBV) infection [13]. Chronic inflammation causes cancer, for instance, in GC through the Correa pathway [14]: chronic gastritis, caused by *H. pylori*, leads to atrophic gastritis, and through a growing number of somatic mutations, progresses from intestinal metaplasia to dysplasia and eventually adenocarcinoma.

The most common subtype in the TCGA classification, chromosomal instability (CIN), is identified by mutated *TP53* and intestinal histology. The microsatellite instability (MSI) subtype is identified by mutations in MSH2, MSH6, PMS2, or MLH1. The EBV-associated subtype is characterized by the *PIK3CA* mutation and *CDKN2A* silencing. The fourth subtype, genetically stable (GS), is associated with a *CDH1* mutation and a diffuse histology, both of which are associated with the loss of cell-to-cell junctions and increased cellular motility. Gastric cancer has been suggested as having immunological phenotypes according to the TCGA classification. For example, the CIN subtype exhibits less T-cell infiltration compared with the EBV subtype, and diffuse GC is associated with increased tertiary lymphoid structures [15].

In this study, we utilized Bio-Rad’s premixed Bio-Plex Pro Human Cytokine 27-plex and 21-plex assay kits to investigate possible serum biomarkers known to play a role in various cancers. We found three serum biomarkers with a prognostic value in GC: C-X-C motif chemokine ligand 12 (CXCL12), also known as stromal-derived factor 1 alpha (SDF-1α), stem cell factor (SCF), also known as the c-KIT ligand named after its receptor c-KIT, and eotaxin, also known as C-C motif chemokine 11 (CCL11).

## Results

### Univariate survival analysis

Multiplex analysis was successful for 29 biomarkers. However, for 18 biomarkers, more than 90% of the observed values fell below the standard curve and were omitted from the final analyses. One biomarker did not yield any results in the multiplex panel. Values for the serum concentrations of the 48 biomarkers were transformed to logarithm base 10 values.

Among the 29 biomarkers, we found three that were statistically significant (*p* < 0.05) using the Cox proportional hazards univariate analysis: CXCL12 (hazard ratio [HR] 0.39, 95% confidence interval [CI] 0.23–0.63, *p* < 0.001, Table 1), SCF (HR 0.38, 95% CI 0.19–0.77, *p* = 0.007), and eotaxin (HR 0.57, 95% CI 0.37–0.89, *p* = 0.013). After FDR correction, CXCL12 and SCF exhibited *p*-values of 0.002 and 0.044, respectively, with low levels indicating a worse survival (Supplementary Table 2).

**Table 1.**
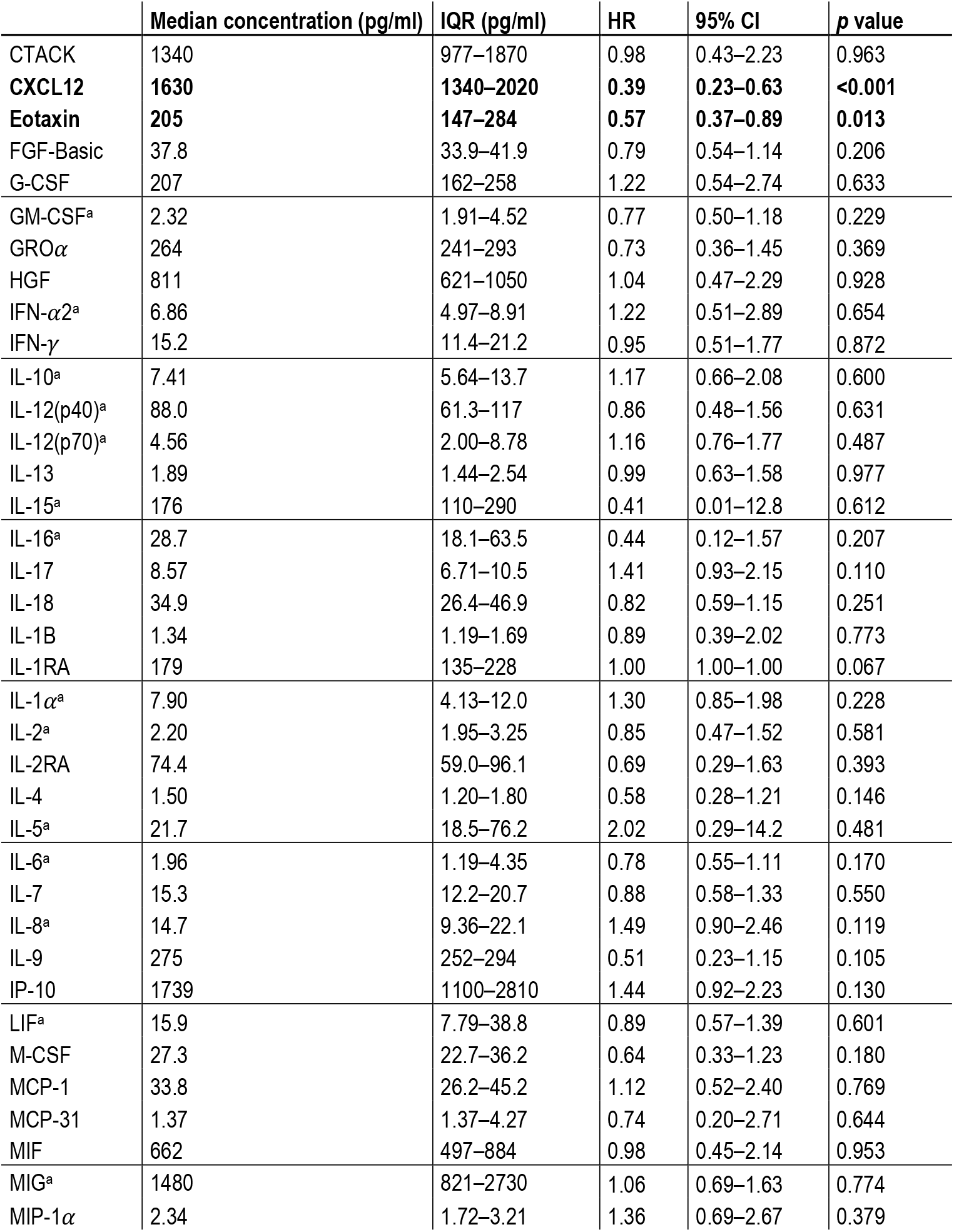
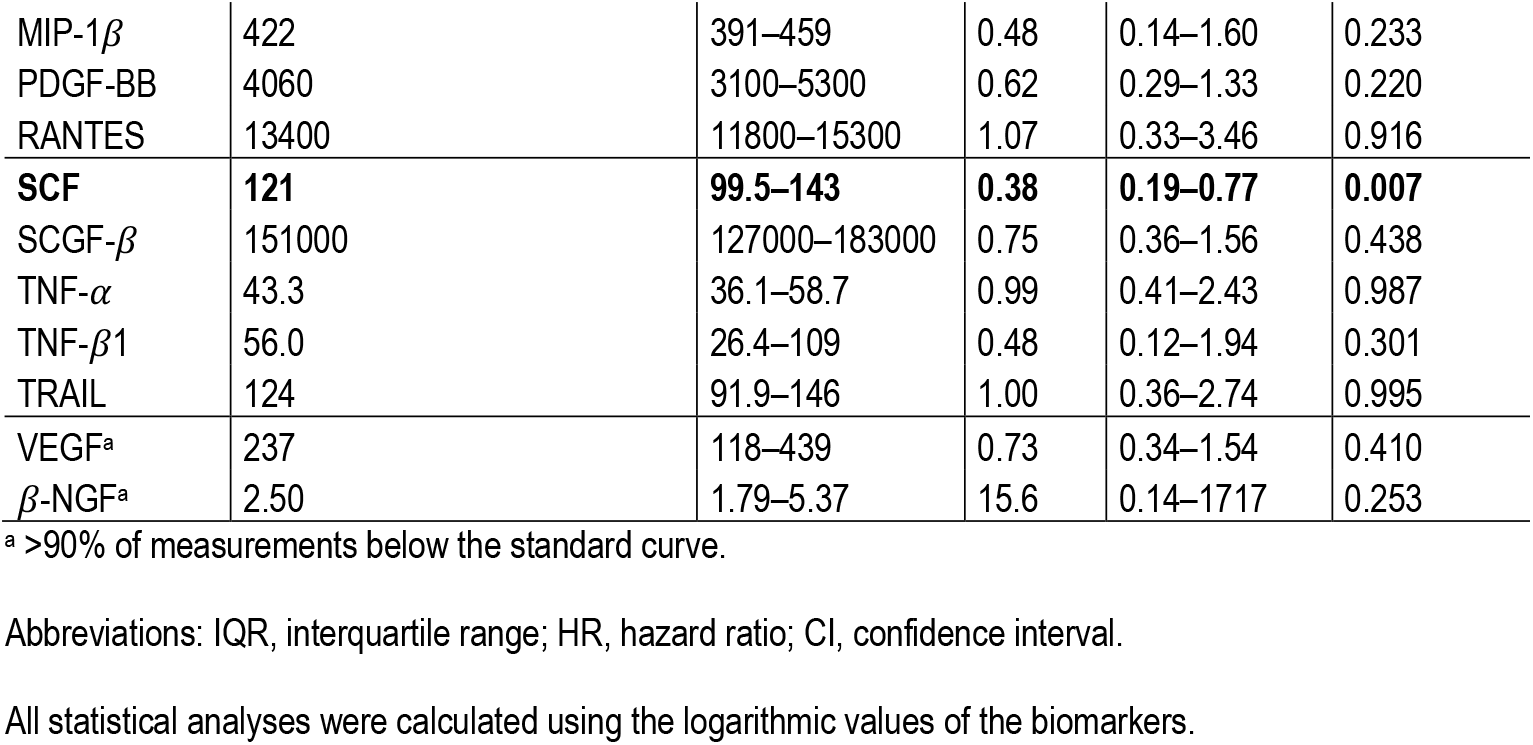
Univariate analysis of biomarkers analyzed using the Bio-Rad’s premixed Bio-Plex Pro Human Cytokine 27-plex assay and 21-plex assay.

**Table 2.** Multivariable Cox regression analysis for disease-specific survival.

|  | **a** |  |  | **b** |  |  | **c** |  |  |
| --- | --- | --- | --- | --- | --- | --- | --- | --- | --- |
|  | **HR** | **95% CI** | **p value** | **HR** | **95% CI** | **p value** | **HR** | **95% CI** | **p value** |
| **Age** |  |  |  |  |  |  |  |  |  |
| <66 | 1.00 |  |  | 1.00 |  |  | 1.00 |  |  |
| ≥66 | 2.63 | 1.82–3.78 | <0.001 | 2.62 | 1.80–3.81 | <0.001 | 2.44 | 1.70–3.50 | <0.001 |
| **Stage** |  |  |  |  |  |  |  |  |  |
| I | 1.00 |  |  | 1.00 |  |  | 1.00 |  |  |
| II | 6.34 | 2.16–18.6 | <0.001 | 6.69 | 2.28–19.6 | <0.001 | 7.45 | 2.54–21.9 | <0.001 |
| III | 22.9 | 8.28–63.3 | <0.001 | 23.3 | 8.43–64.4 | <0.001 | 25.2 | 9.10–70.0 | <0.001 |
| IV | 75.1 | 25.9–218 | <0.001 | 72.9 | 25.2–211 | <0.001 | 83.3 | 28.7–242 | <0.001 |
| **Laurén classification** |  |  |  |  |  |  |  |  |  |
| Intestinal | 1.00 |  |  | 1.00 |  |  | 1.00 |  |  |
| Diffuse and mixed | 2.45 | 1.65–3.65 | <0.001 | 2.17 | 1.47–3.21 | <0.001 | 2.25 | 1.52–3.34 | <0.001 |
| **CXCL12** | 0.13 | 0.04–0.48 | 0.002 |  |  |  |  |  |  |
| **SCF** |  |  |  | 0.42 | 0.11–1.57 | 0.196 |  |  |  |
| **Eotaxin** |  |  |  |  |  |  | 0.40 | 0.19–0.86 | 0.018 |

Abbreviations: HR, hazard ratio; CI, confidence interval; CXCL12, C-X-C motif chemokine ligand 12; SCF, stem cell factor.
115 All statistical analyses were calculated using the logarithmic values of the biomarkers.

### Multivariate survival analysis

In a multivariate analysis adjusted for age, stage, and the histological Laurén classification, CXCL12 and eotaxin emerged as statistically significant: CXCL12 had a HR of 0.13 (95% CI 0.04–0.49, *p* = 0.002, Table 2a), SCF had a HR of 0.42 (95% CI 0.11–1.57, *p* = 0.196; Table 2b), and eotaxin had a HR of 0.40 (95% CI 0.19–0.86, *p* = 0.018; Table 2c).

### Survival analysis of CXCL12, SCF, and eotaxin in patient subgroups

The ability to predict disease-specific survival was analyzed using the time-dependent AUC values with 95% confidence intervals (Figure 1). Of the three biomarkers, CXCL12 was statistically significant at the majority of time points (Figure 1a). In the ROC curve diagrams, at the 10-year time point CXCL12 had an AUC of 63.9% (95% CI 52.8–74.9). The three biomarkers were dichotomized based on the maximum value of Youden’s index for the Kaplan–Meier analysis [16].

**Fig. 1.**
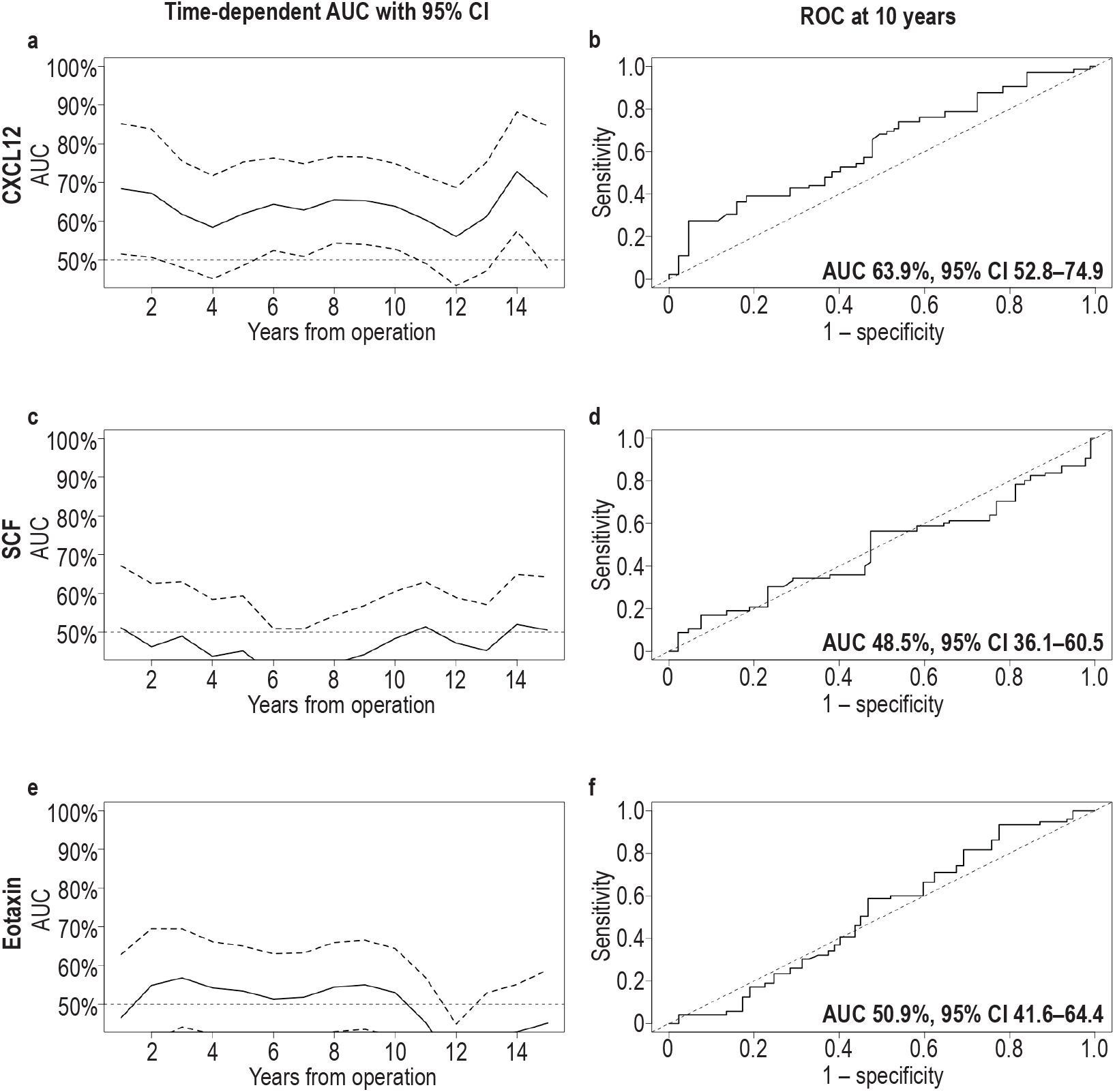
Time-dependent analysis of the area under the curve (AUC) and receiver operating characteristic (ROC) curve at the 10-year time point with 95% confidence intervals for CXCL12 (a and b), SCF (c and d), and eotaxin (e and f), respectively. Abbreviations: AUC, area under the curve; ROC, receiver operating characteristic; CI, confidence interval; CXCL12, C-X-C motif chemokine ligand 12; SCF, stem cell factor.

The estimated cutoff point for CXCL12 was 1513 pg/ml. The 5-year DSS among patients with high serum levels was 53.2% (95% CI 45.3–62.5), falling to 32.1% (95% CI 23.6–43.5, log rank test: *p* < 0.001, Figure 2a) with low serum CXCL12 levels. The cutoff point for SCF was 97 pg/ml. The 5-year DSS among patients with high serum SCF levels was 48.1% (95% CI 41.2–56.2), falling to 33.6% (95% CI 22.1–51.0, *p* = 0.073, Figure 2b) with low serum levels. The cutoff point for eotaxin was 267 pg/ml. The 5-year DSS among patients with high serum eotaxin levels was 54.4% (95% CI 43.4– 68.2), falling to 41.3% (95% CI 34.2–49.9, *p* = 0.037, Figure 2c) with low serum levels.

**Fig. 2.**
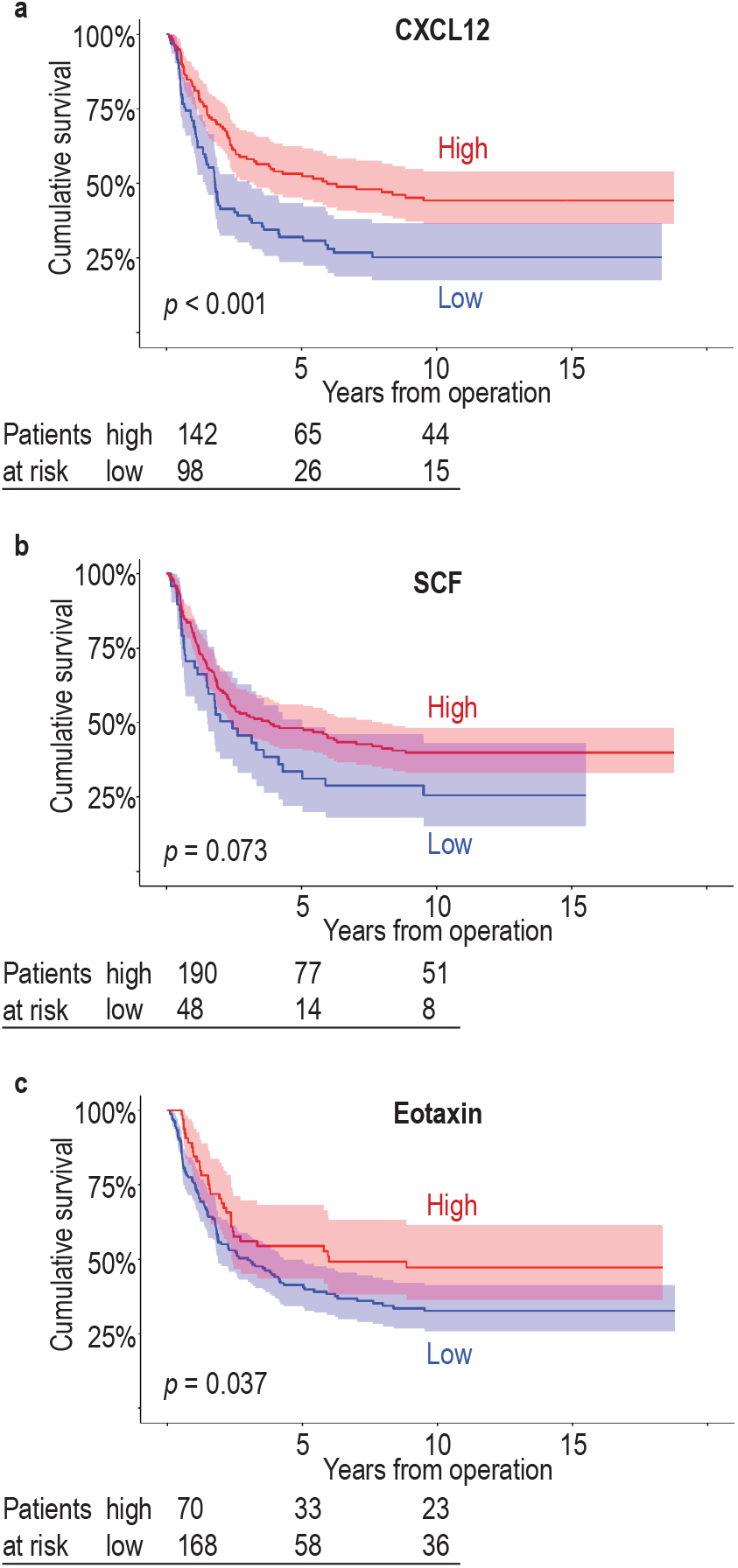
Disease-specific survival of patients with either a high or low serum concentration of CXCL12 (a), SCF (b), and eotaxin (c). Survival curves were drawn according to the Kaplan–Meier method and the *p*-values were calculated using the log-rank test. Abbreviations: CXCL12, C-X-C motif chemokine ligand 12; SCF, stem cell factor.

Among patients with a diffuse histology, the three serum biomarkers, CXCL12 (*p* < 0.001; Figure 3a and Supplementary Table 3), SCF (*p* = 0.010; Figure 3d), and eotaxin (*p* = 0.022; Figure 3g), all served as prognostic factors. Among patients with lymph node metastases (pN+), those with higher levels of CXCL12 (*p* < 0.001; Figure 3b) and eotaxin (*p* = 0.026; Figure 3h) exhibited a better survival.

**Table 3.** Associations between serum biomarker levels and clinicopathological variables.

|  | **CXCL12<sup>high</sup>**<br>n = 142 (59.2%) | **p value<sup>a</sup>** | **SCF<sup>high</sup>**<br>n = 190 (79.8%) | **p value<sup>a</sup>** | **Eotaxin<sup>high</sup>**<br>n = 70 (29.5%) | **p value<sup>a</sup>** |
| --- | --- | --- | --- | --- | --- | --- |
| **Age** |  |  |  |  |  |  |
| <66 | 75 (61.0) | 0.559 | 87 (71.3) | **<0.001** | 37 (30.3) | 0.783 |
| ≥66 | 67 (57.3) |  | 103 (88.8) |  | 33 (28.7) |  |
| **Sex** |  |  |  |  |  |  |
| Male | 70 (59.8) | 0.839 | 96 (82.1) | 0.423 | 39 (33.3) | 0.192 |
| Female | 72 (58.5) |  | 94 (77.7) |  | 31 (25.6) |  |
| **Stage** |  |  |  |  |  |  |
| I | 36 (73.5) | 0.078<sup>b</sup> | 44 (89.8) | **0.005<sup>b</sup>** | 15 (30.6) | 0.281<sup>b</sup> |
| II | 29 (54.7) |  | 46 (86.8) |  | 17 (31.5) |  |
| III | 54 (56.3) |  | 71 (75.5) |  | 31 (33.3) |  |
| IV | 23 (54.8) |  | 29 (69.0) |  | 7 (16.7) |  |
| **Tumor invasion (pT)** |  |  |  |  |  |  |
| 1 | 27 (75.0) | 0.101<sup>b</sup> | 32 (88.9) | 0.066<sup>b</sup> | 14 (38.9) | 0.379<sup>b</sup> |
| 2 | 22 (61.1) |  | 32 (88.9) |  | 10 (27.8) |  |
| 3 | 42 (52.5) |  | 59 (74.7) |  | 21 (26.3) |  |
| 4 | 51 (58.0) |  | 67 (77.0) |  | 25 (29.1) |  |
| **Lymph node metastasis (pN)** |  |  |  |  |  |  |
| No | 47 (63.5) | 0.444 | 63 (85.1) | 0.165 | 25 (33.3) | 0.384 |
| Yes | 92 (58.2) |  | 120 (76.9) |  | 43 (27.7) |  |
| **Distant metastasis (M)** |  |  |  |  |  |  |
| No | 119 (60.1) | 0.523 | 161 (82.1) | 0.060 | 63 (32.1) | **0.046** |
| Yes | 23 (54.8) |  | 29 (69.0) |  | 7 (16.7) |  |
| **Laurén classification** |  |  |  |  |  |  |
| Intestinal | 47 (54.7) | 0.288 | 68 (80.0) | 0.962 | 31 (36.9) | 0.061 |
| Diffuse and other | 95 (61.7) |  | 122 (79.7) |  | 39 (25.3) |  |
| **MMR** |  |  |  |  |  |  |
| MMRp | 106 (61.6) | 0.440 | 135 (78.9) | 0.883 | 52 (30.4) | 0.499 |
| MMRd | 22 (55.0) |  | 32 (80.0) |  | 10 (25.0) |  |
| **EBV *ish*** |  |  |  |  |  |  |
| EBV negative | 127 (60.5) | 0.908 | 166 (79.4) | 0.577 | 58 (27.8) | **0.034** |
| EBV positive | 5 (62.5) |  | 7 (87.5) |  | 5 (62.5) |  |
| **p53 staining** |  |  |  |  |  |  |
| Aberrant | 105 (62.9) | 0.230 | 134 (80.7) | 0.350 | 48 (28.9) | 0.908 |
| Wild type | 25 (53.2) |  | 35 (74.5) |  | 14 (29.8) |  |

<sup>a</sup>Pearson chi-square. <sup>b</sup>Linear-by-linear association.
Abbreviations: CXCL12, C-X-C motif chemokine ligand 12; SCF, stem cell factor; MMRp/d, mismatch repair proficient/deficient; EBVish, Epstein–Barr virus *in situ* hybridization.

**Fig. 3.**
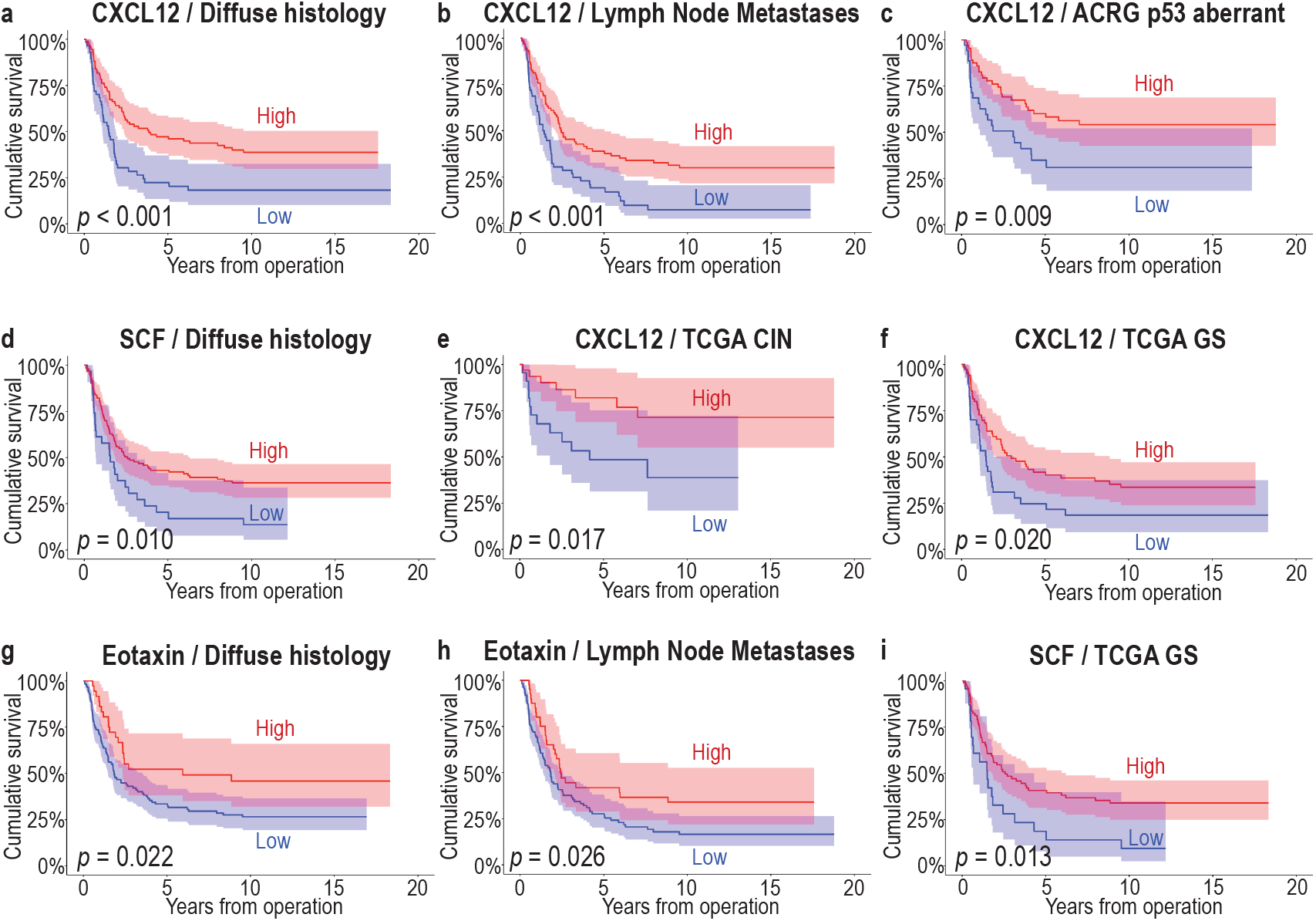
Disease-specific survival of patients with either a high or low serum concentration of CXCL12, SCF, and eotaxin in different patient subgroups. Patients with diffuse histology (a), lymph node metastasis (b), or belonging to the p53 aberrant group in the ACRG classification (c), CIN (e) or GS (f) groups in the TCGA classification exhibited a better survival when CXCL12 levels were higher. Patients with a diffuse histology (d) or patients belonging to the GS group (i) of the TCGA classification experienced a better survival when the SCF levels were higher. In addition, in patients with high eotaxin levels, patients with a diffuse histology (g) or lymph node metastasis (h) exhibited a better survival. Survival curves were drawn according to the Kaplan–Meier method and the *p*-values were calculated using the log-rank test. Abbreviations: ACRG, Asian Cancer Research Group; CIN, chromosomal instability; CXCL12, C-X-C motif chemokine ligand 12; GS, genetically stable; SCF, stem cell factor; TCGA, The Cancer Genome Atlas.

We previously identified immunohistochemically determined prognostic patient subgroups using classifications by TCGA and ACRG [17]. Patients with high CXCL12 levels exhibited a better prognosis in the immunohistochemical ACRG classification’s p53 aberrant subtype (*p* = 0.009; Figure 3c), with the TCGA classification’s subtypes chromosomal instability (CIN) characterized by an intestinal histology (*p* = 0.017; Figure 3e) and genetically stable (GS) identified by a diffuse histology (*p* = 0.020; Figure 3f). Similarly, patients with a high SCF concentration exhibited a better survival in the immunohistochemical TCGA classification’s GS subtype (*p* = 0.013; Figure 3i).

### Associations between serum concentrations and clinicopathological variables

A high serum concentration of CXCL12 did not associate with the clinicopathological variables we examined (Table 3). However, a high serum concentration of SCF associated with an older age (*p* < 0.001) and stage of disease (*p* = 0.005), and a high serum concentration of eotaxin associated with distant metastasis (*p* = 0.046) and EBV positivity (*p* = 0.034).

## Discussion

Multivariate survival analysis revealed that, among the 48 biomarkers we analyzed, CXCL12 and eotaxin served as independent prognostic markers among gastric cancer patients. Additionally, high levels of SCF are also biomarkers indicative of a better survival. Subgroup analysis further revealed new associations between these novel serum biomarkers and other prognostic markers, similar to the Laurén classification and molecular subtypes described in our previous work [17].

CXCL12 is a well-known cytokine associated with many pathologies and malignancies via signaling with its receptors CXCR4 and CXCR7 [18]. In cancer, CXCL12 is involved in tumor progression, angiogenesis, metastasis, and survival through various downstream signaling pathways [18]. We found that, contrary to other studies [19, 20], high levels of CXCL12 serve as independent marker for a better prognosis in GC. The serum levels of CXCL12 have been studied less than the expression in tumor tissue. Tumor and serum levels of CXCL12 are not necessarily comparable. CXCL12 can be expressed on the cell surface of both cancer cells as well as immune and other stromal cells [18]. Furthermore, CXCL12 can be either secreted into the bloodstream, locally in the tissue, or just expressed on the cell surface. Our results concerning the serum levels of CXCL12 do not take into consideration which cells have secreted it nor how it could be expressed in the tissue. A meta-analysis of CXCL12 expression showed that patients with high CXCL12 levels in the serum or tumor experienced a worse survival in esophagogastric and pancreatic cancer, whereas, in ovarian and colorectal cancer, the effect remained unclear [19]. In another study, gastric cancer patients with a high expression of CXCL12 in the tumor tissue did not indicate a better survival [20]. The AUC value of CXCL12 in our study was a moderate 63.9%, but was statistically significant at a majority of the time points analyzed, suggesting its ability to effectively differentiate between groups. More recently, the role of CXCL12 in the tumor microenvironment (TME) and especially cancer-associated fibroblasts (CAF) have been further explored. A higher CXCL12 expression associated with a pro-invasive inflammatory subset of CAFs, leading to worse clinical outcomes [21]. In the subgroup analysis, we observed patients with a diffuse histology or lymph node metastasis still exhibiting a better prognosis when they had higher serum levels of CXCL12. The serum and tumor levels of CXCL12 previously associated with lymph node and distant metastases of GC [22, 23]. However, we found no association between elevated serum levels of CXCL12 and the clinical or histological variables. Our data suggest that high CXCL12 levels in the serum are a systemic marker for a better survival.

The stem cell factor (SCF) is primarily expressed in hematopoietic cells, but can also be found in the gastric epithelium’s peristaltic pacemaker cells called interstitial cells of Cajal (ICCs) [24]. The tissue expression of SCF and its receptor c-KIT has been previously identified in GC [25, 26], although their prognostic effect has not been previously identified. We found that patients with high serum levels of SCF exhibited a better prognosis, although we did not identify a single cutoff point to categorize patients into distinct groups of better and worse survival. The serum levels of SCF have been associated with GC in a multiplex setting alongside 18 other proteins as a proteomic tool for GC diagnosis, but not as an independent biomarker [27]. The c-KIT mutation can be found in different types of cancer, but one of the most studied examples of c-KIT’s involvement in cancer is the gastrointestinal stromal tumor (GIST), which originates from ICCs [24]. Patients with GIST have elevated tissue levels of SCF and advanced disease presumably by autocrine and paracrine activation of c-KIT signaling [28]. However, in another study, the serum levels of SCF were lower in patients with GIST compared with healthy controls [29]. One study suggests that *H. pylori* infection causes the downregulation of SCF expression in gastric tissue leading to fewer ICCs [30]. Injections of SCF have been used as a treatment for metastatic GC in a mouse model, where a subsequent increased number in and activity of ICCs promoted normal peristaltic activity [31]. Low serum levels of SCF may indicate other non-beneficial effects on patients such as an impaired function of the gastrointestinal tract, whereas high serum levels promote normal function. We found an association between high SCF serum levels and a lower stage of disease, yet no association with lymph node or distant metastasis. We also found that patients with low SCF levels exhibited a worse prognosis.

Eotaxin is a chemokine primarily involved in guiding eosinophils, thus primarily affecting allergic reactions [32]. We identified eotaxin as a novel independent prognostic biomarker in GC. While the prognostic effect of eotaxin had previously appeared unclear, EBV-positive GC had associated with elevated plasma levels of eotaxin, confirmed by our results as well [33]. In another study, no associations between eotaxin levels and clinicopathological variables were observed [34].

The elevated presence of eotaxin was noted in different malignancies: colorectal cancer, oral squamous cell cancer, and breast cancer [32]. Angiogenesis and evading apoptosis were previously identified roles played by eotaxin, which may also play a part in GC [35, 36].

We previously identified prognostic serum biomarkers using the same multiplex panel of 48 serum biomarkers exploring levels of cytokines and growth factors in colorectal cancer and pancreatic cancer [37, 38]. In one previous study, the prognostic effect of serum biomarkers was explored, resulting in four candidates for biomarkers of worse survival in GC: IL-10Rb, adenosine deaminase (ADA), IL-20, and oncostatin M (OSM), which were not included in our panel of 48 cytokines [39]. Additionally, multiplex analysis was used to identify diagnostic marker combinations for GC [40, 41]. These findings, however, have not shed any further light on improving GC care. Preliminary results demand further validation before consideration for clinical use, preferably in different patient cohorts.

In our study, we used a rather large patient cohort of 240 patients comprised almost entirely of gastric cancer patients undergoing surgery at Helsinki University Hospital within a specific time frame. Yet, a larger cohort would be needed for the further validation of our results. It is also worth noting that most serum samples were frozen at –80°C for over ten years before first thawing for our analysis. It is unknown how the proteins tested may change resulting from long-time storage. Our previous studies using samples stored for prolonged time periods have been successful [37, 38, 42, 43]. However, this was the first time the samples were thawed, which is likely the most crucial point in preserving samples for longer times. This has also allowed for a very long follow-up period of over ten years. A considerable number of the biomarkers examined were omitted from further analysis due to too many values falling outside the range of control values. One reason behind the higher number of excluded cytokines here than in our previous analyses [37, 38, 42, 43] could be the variation between different manufacturing lots of the multiplex panel kit used. Other benefits of an older patient cohort are that there are almost no neoadjuvant treated patients, making it possible to have comprehensive clinical data combined with histological samples without the possible influence of neoadjuvant chemotherapy.

Both CXCL12 and eotaxin as chemokines primarily associated with inducing inflammation further underline the association between GC and the infectious agents behind it. Proinflammatory molecules may be indicative of a stronger immune response against tumors, which could partly explain why patients with a higher systemic expression of these biomarkers experience a better prognosis. Our results point to the need to further examine the immunological landscapes of gastric cancer as possible new targets for prognostic evaluation and maybe even treatment.

## Conclusions

The serum biomarkers CXCL12, SCF, and eotaxin can be used to assess the prognosis for gastric cancer patients. The prognostic effect of inflammatory serum biomarkers CXCL12 and eotaxin in gastric cancer might shed new light on our understanding of the immunological microenvironment of gastric cancer. Further investigation of these findings might yield new possibilities for treatments and help to better target treatments to specific patient subgroups.

## Materials and methods

### Patients

The patient cohort comprised 240 individuals who underwent surgery for histologically verified gastric adenocarcinoma in the Department of Surgery, Helsinki University Hospital, between 2000 and 2009. We excluded patients with a history of malignant disease or synchronous cancer. The median age of the patient cohort at the time of surgery was 65.6 years (interquartile range [IQR] 56.5–75.5; Supplementary Table 1). Among all patients, 117 (48.8%) were male, and the median survival was 2.29 years (IQR 0.90–9.94). For staging, we used the seventh version of the tumor-node-metastasis (TNM) classification [44]. Overall, 49 (20.4%) had stage I cancer, 53 (22.1%) had stage II, 96 (40.0%) had stage III, and 42 (17.5%) had stage IV. An intestinal histology according to the Laurén classification was observed in 86 patients (35.8%), while 154 (64.2%) had diffuse or other type of histology. Adjuvant chemotherapy was administered to 100 patients (43.7%), and 43 patients (19.1%) received adjuvant radiotherapy. Only 13 patients (5.4%) received neoadjuvant chemotherapy.

### Serum samples

Serum samples were obtained from patients after possible neoadjuvant therapy, shortly before surgery, aliquoted, and subsequently stored at –80°C until the multiplex assay was performed in 2018.

### Protein profiling

To determine the serum protein concentrations of cytokines and growth factors, we used Bio-Rad’s premixed Bio-Plex Pro Human Cytokine 27-plex assay (catalog no. M500KCAF0Y) and 21-plex assay (catalog no. MF0005KMII) kits on Bio-Rad’s Bio-Plex 200 system (Supplementary Table 2). Assays were used according to the manufacturer’s instructions. However, only half the recommended concentration levels were used to detect the number of antibodies, beads, and the streptavidin–phycoerythrin conjugate. We validated this approach in our previous studies [42, 43], and the approach was used successfully in other cancer patient cohorts [37, 38].

### Immunohistochemistry and determining the phenotypic subtypes

We constructed a tumor tissue microarray immunostained for the following markers: MSI markers MSH2, MSH6, MLH1, and PMS2; p53; E-cadherin; and EBER*ISH*. We used these stainings to divide patients into phenotypic subtypes according to the molecular subtypes of the TCGA and ACRG classifications [17].

## Statistical analysis

We used two-tailed *p*-values and considered *p* < 0.05 as statistically significant. Statistical evaluations were calculated using IBM’s statistical software (IBM SPSS Statistics Version 28, International Business Machines Corp., NY, USA) and R (R version 4.3.1, Foundation for Statistical Computing, Vienna, Austria). Associations between groups and continuous variables were assessed using the Mann–Whitney U-test and the Kruskal–Wallis test. For the univariate and multivariate analyses, we used the Cox proportional hazards regression analysis to calculate the disease-specific survival (DSS), and applied the false discovery rate (FDR) correction for multiple testing [45]. We defined DSS as the time from surgery until death from GC or until the end of the follow-up period. We chose patient characteristics consisting of patient age, stage (categorical variable), and the Laurén classification for the multivariate survival analysis using the Cox regression model. We calculated the time-dependent receiver operating characteristic (ROC) curves and the area under the curves (AUCs) using the TimeROC package in R, and the integrated AUC over time from 1 to 15 years. For the dichotomization of the biomarkers, we used the maximum value of Youden’s index at the 10-year time point [16]. For figures with Kaplan–Meier curves, statistical significance was calculated using the log-rank analysis.

## Ethics approval

The research conducted was approved by the appropriate authorities: Finnish National Supervisory Authority of Health and Welfare (permit number for research conducted with human samples: Valvira Dnro 1004/06.01.03.01/2012), the hospital district of Helsinki and Uusimaa (permit number HUS/23/2024), and the Ethics Committee of Medicine of Helsinki University Hospital (permit number HUS/1223/2021). Patient information, samples, and data were handled and stored in accordance with research permits, the Declaration of Helsinki and other local regulations and guidelines.

## Consent to participate

Patients provided their written informed consent to participate in the study prior to giving blood samples.

## Acknowledgements

We thank Pia Saarinen and Maria Finne for their essential technical assistance, and Vanessa Fuller for English-language revisions.

## Author contributions

Brodkin, Kaprio, Haglund, and Böckelman contributed to the study conception and design. Material preparation and data collection were performed by Leppä, Kokkola, Salmi, and Jalkanen. Statistical analysis and interpretation were conducted by Brodkin, Kaprio, Mustonen, and Böckelman. The first draft of the manuscript was written by Brodkin, Kaprio, and Böckelman, and all authors commented on the manuscript. All authors read and approved the final manuscript.

## Data availability

The datasets supporting the conclusions of this article are included within the article and its supplementary files. Other data used in this study are available from the corresponding author upon reasonable request.

## Abbreviations

ACRG: Asian Cancer Research Group
AUC: area under the curve
CAF: cancer-associated fibroblasts
CA19-9: carbohydrate antigen 19-9
CCL11: C-C motif chemokine 11
CEA: carcinoembryonic antigen
CI: confidence interval
CIN: chromosomal instability
CRP: C-reactive protein
CXCL12: C-X-C motif chemokine ligand 12
DSS: disease-specific survival
EBV: Epstein–Barr virus
FDR: false discovery rate
GC: gastric cancer
GS: genetically stable
HR: hazard ratio
ICC: interstitial cells of Cajal
IHC: immunohistochemistry
IQR: interquartile range
MMRp: mismatch repair proficiency
OS: overall survival
ROC: receiver operating characteristic
SCF: stem cell factor
SDF-1: stromal cell-derived factor 1 alpha
TCGA: The Cancer Genome Atlas
TME: tumor microenvironment
TNM: tumor-node-metastasis classification

