## supplementary data for "CXCL12, SCF, and eotaxin are prognostic serum biomarkers in gastric cancer"

Supplementary Table 1. Patient characteristics

| | n = 240 (%) |
|---|---|
| **Age** | |
| <66 | 123 (51.2) |
| ≥66 | 117 (48.8) |
| **Sex** | |
| Male | 117 (48.8) |
| Female | 123 (51.2) |
| **Stage** | |
| I | 49 (20.4) |
| II | 53 (22.1) |
| III | 96 (40.0) |
| IV | 42 (17.5) |
| **Tumor invasion (pT)** | |
| 1 | 36 (15.0) |
| 2 | 36 (15.0) |
| 3 | 80 (33.3) |
| 4 | 88 (36.7) |
| **Lymph node metastasis (pN)** | |
| No | 74 (31.9) |
| Yes | 158 (68.1) |
| **Distant metastasis (M)** | |
| No | 198 (82.5) |
| Yes | 42 (17.5) |
| **Laurén classification** | |
| Intestinal | 86 (35.8) |
| Diffuse and other | 154 (64.2) |
| **Adjuvant chemotherapy** | |
| No | 129 (56.3) |
| Yes | 100 (43.7) |
| **Adjuvant radiotherapy** | |
| No | 182 (80.9) |
| Yes | 43 (19.1) |
| **Neoadjuvant chemotherapy** | |
| No | 227 (94.6) |
| Yes | 13 (5.4) |

Supplementary Table 2. Univariate analysis of biomarkers analyzed using Bio-Rad's premixed Bio-Plex Pro Human Cytokine 27-plex and 21-plex assays

| | | HR | 95% CI | *p* value | FDR-corrected *p* value |
|---|---|---|---|---|---|
| CTACK | Cutaneous T cell-attracting chemokine | 0.98 | 0.43–2.23 | 0.963 | 0.995 |
| **CXCL12** | **C-X-C motif chemokine ligand 12** | **0.39** | **0.23–0.63** | **<0.001** | **0.002** |
| **Eotaxin** | | **0.57** | **0.37–0.89** | **0.013** | **0.066** |
| FGF-Basic | Basic fibroblast growth factor | 0.79 | 0.54–1.14 | 0.206 | 0.494 |
| G-CSF | Granulocyte colony-stimulating factor | 1.22 | 0.54–2.74 | 0.633 | 0.876 |
| GM-CSF[a] | Granulocyte-macrophage colony-stimulating factor | 0.77 | 0.50–1.18 | 0.229 | |
| GRO$\alpha$ | Growth-regulated oncogene alpha | 0.73 | 0.36–1.45 | 0.369 | 0.673 |
| HGF | Hepatocyte growth factor | 1.04 | 0.47–2.29 | 0.928 | 0.995 |
| IFN- 2[a] | Interferon alpha 2 | 1.22 | 0.51–2.89 | 0.654 | |
| IFN- | Interferon gamma | 0.95 | 0.51–1.77 | 0.872 | 0.995 |
| IL-10[a] | Interleukin 10 | 1.17 | 0.66–2.08 | 0.600 | |
| IL-12(p40)[a] | Interleukin 12 (p40) | 0.86 | 0.48–1.56 | 0.631 | |
| IL-12(p70)[a] | Interleukin 12 (p70) | 1.16 | 0.76–1.77 | 0.487 | |
| IL-13 | Interleukin 13 | 0.99 | 0.63–1.58 | 0.977 | 0.995 |
| IL-15[a] | Interleukin 15 | 0.41 | 0.01–12.8 | 0.612 | |
| IL-16[a] | Interleukin 16 | 0.44 | 0.12–1.57 | 0.207 | |
| IL-17 | Interleukin 17 | 1.41 | 0.93–2.15 | 0.110 | 0.395 |
| IL-18 | Interleukin 18 | 0.82 | 0.59–1.15 | 0.251 | 0.502 |
| IL-1RA | Interleukin 1 receptor antagonist | 1.00 | 1.00–1.00 | 0.067 | 0.303 |
| IL-1 [a] | Interleukin 1 alpha | 1.30 | 0.85–1.98 | 0.228 | |
| IL-1$\beta$ | Interleukin 1 beta | 0.89 | 0.39–2.02 | 0.773 | 0.994 |
| IL-2[a] | Interleukin 2 | 0.85 | 0.47–1.52 | 0.581 | |
| IL-2RA | Interleukin 2 receptor antagonist | 0.69 | 0.29–1.63 | 0.393 | 0.673 |
| IL-4 | Interleukin 4 | 0.58 | 0.28–1.21 | 0.146 | 0.439 |
| IL-5[a] | Interleukin 5 | 2.02 | 0.29–14.2 | 0.481 | |
| IL-6[a] | Interleukin 6 | 0.78 | 0.55–1.11 | 0.170 | |
| IL-7 | Interleukin 7 | 0.88 | 0.58–1.33 | 0.550 | 0.826 |
| IL-8[a] | Interleukin 8 | 1.49 | 0.90–2.46 | 0.119 | |
| IL-9 | Interleukin 9 | 0.51 | 0.23–1.15 | 0.105 | 0.395 |
| IP-10 | Interferon gamma-induced protein 10 | 1.44 | 0.92–2.23 | 0.130 | 0.427 |
| LIF[a] | Leukemia inhibitory factor | 0.89 | 0.57–1.39 | 0.601 | |
| M-CSF | Macrophage colony-stimulating factor | 0.64 | 0.33–1.23 | 0.180 | 0.494 |
| MCP-1 | Monocyte chemoattractant protein 1 | 1.12 | 0.52–2.40 | 0.769 | 0.994 |
| MCP-3[a] | Monocyte chemoattractant protein 3 | 0.74 | 0.20–2.71 | 0.644 | |
| MIF | Macrophage migration inhibitory factor | 0.98 | 0.45–2.14 | 0.953 | 0.995 |
| MIG[a] | Monokine induced by gamma interferon | 1.06 | 0.69–1.63 | 0.774 | |
| MIP-1$\alpha$ | Macrophage inflammatory protein 1 alpha | 1.36 | 0.69–2.67 | 0.379 | 0.673 |
| MIP-1$\beta$ | Macrophage inflammatory protein 1 beta | 0.48 | 0.14–1.60 | 0.233 | 0.494 |
| PDGF-BB | Platelet-derived growth factor BB | 0.62 | 0.29–1.33 | 0.220 | 0.494 |
| RANTES | Regulated on activation, normal T cell expressed and secreted | 1.07 | 0.33–3.46 | 0.916 | 0.995 |
| **SCF** | **Stem cell factor** | **0.38** | **0.19–0.77** | **0.007** | **0.044** |
| SCGF-$\beta$ | Serum stem cell growth factor beta | 0.75 | 0.36–1.56 | 0.438 | 0.716 |
| TNF-$\alpha$ | Tumor necrosis factor alpha | 0.99 | 0.41–2.43 | 0.987 | 0.995 |
| TNF- [a] | Tumor necrosis factor beta | 0.48 | 0.12–1.94 | 0.301 | |
| TRAIL | TNF-related apoptosis-inducing ligand | 1.00 | 0.36–2.74 | 0.995 | 0.995 |
| VEGF[a] | Vascular endothelial growth factor | 0.73 | 0.34–1.54 | 0.410 | |
| -NGF[a] | Nerve growth factor beta | 15.6 | 0.14–1717 | 0.253 | |

[a]Measurements fell beyond the standard curve.

Abbreviations: IQR, interquartile range; HR, hazard ratio; CI, confidence interval; FDR, false discovery rate.

All statistical analyses were completed using the logarithmic values of the biomarkers.

Supplementary Table 3. Univariate survival analysis of serum biomarkers where low serum levels serve as the reference value (HR = 1.00)

| | CXCL12 | | | SCF | | | Eotaxin | | |
|---|---|---|---|---|---|---|---|---|---|
| | HR | 95% CI | *p* value | HR | 95% CI | *p* value | HR | 95% CI | *p* value |
| **Age** | | | | | | | | | |
| <66 | 0.44 | 0.27–0.70 | **<0.001** | 0.42 | 0.26–0.69 | **<0.001** | 0.63 | 0.36–1.08 | 0.094 |
| ≥66 | 0.74 | 0.46–1.20 | 0.216 | 1.60 | 0.69–3.70 | 0.277 | 0.72 | 0.41–1.26 | 0.244 |
| **Sex** | | | | | | | | | |
| Male | 0.64 | 0.39–1.06 | 0.085 | 0.73 | 0.39–1.34 | 0.309 | 0.82 | 0.48–1.40 | 0.474 |
| Female | 0.49 | 0.31–0.78 | **0.002** | 0.69 | 0.41–1.15 | 0.151 | 0.52 | 0.29–0.95 | **0.034** |
| **Stage** | | | | | | | | | |
| I | 1.14 | 0.12–11.0 | 0.910 | N/A | | | 0.81 | 0.08–7.78 | 0.855 |
| II | 0.55 | 0.23–1.31 | 0.176 | 1.32 | 0.31–5.69 | 0.711 | 0.39 | 0.13–1.15 | 0.088 |
| III | 0.46 | 0.29–0.73 | **<0.001** | 1.04 | 0.62–1.76 | 0.878 | 0.86 | 0.52–1.41 | 0.541 |
| IV | 1.00 | 0.53–1.92 | 0.989 | 0.91 | 0.45–1.81 | 0.781 | 0.40 | 0.16–1.04 | 0.061 |
| **Tumor invasion (pT)** | | | | | | | | | |
| 1 | 0.16 | 0.02–1.78 | 0.137 | N/A | | | 0.79 | 0.07–8.69 | 0.845 |
| 2 | 0.89 | 0.26–3.01 | 0.855 | 0.52 | 0.11–2.43 | 0.409 | 0.86 | 0.23–3.25 | 0.821 |
| 3 | 0.56 | 0.33–0.96 | **0.034** | 1.05 | 0.56–1.97 | 0.871 | 0.81 | 0.43–1.50 | 0.497 |
| 4 | 0.65 | 0.41–1.05 | 0.075 | 0.76 | 0.44–1.30 | 0.307 | 0.53 | 0.30–0.93 | **0.027** |
| **Lymph node metastasis (pN)** | | | | | | | | | |
| No | 0.59 | 0.23–1.54 | 0.281 | 0.55 | 0.18–1.69 | 0.297 | 0.82 | 0.29–2.31 | 0.712 |
| Yes | 0.51 | 0.35–0.74 | **<0.001** | 0.79 | 0.52–1.22 | 0.289 | 0.61 | 0.39–0.95 | **0.027** |
| **Distant metastasis (M)** | | | | | | | | | |
| No | 0.48 | 0.32–0.71 | **<0.001** | 0.77 | 0.47–1.24 | 0.278 | 0.80 | 0.52–1.24 | 0.324 |
| Yes | 1.00 | 0.53–1.92 | 0.989 | 0.91 | 0.45–1.81 | 0.781 | 0.40 | 0.16–1.04 | 0.061 |
| **Laurén classification** | | | | | | | | | |
| Intestinal | 0.54 | 0.29–1.02 | 0.058 | 1.18 | 0.52–2.67 | 0.700 | 1.01 | 0.52–1.96 | 0.981 |
| Diffuse and other | 0.51 | 0.34–0.76 | **<0.001** | 0.56 | 0.38–0.88 | **0.012** | 0.56 | 0.34–0.93 | **0.024** |
| **MMR** | | | | | | | | | |
| MMRp | 0.56 | 0.38–0.83 | **0.004** | 0.67 | 0.43–1.06 | 0.085 | 0.73 | 0.46–1.14 | 0.163 |
| MMRd | 0.77 | 0.36–1.67 | 0.508 | 0.95 | 0.38–2.37 | 0.911 | 0.43 | 0.15–1.24 | 0.119 |
| **EBV *ish*** | | | | | | | | | |
| EBV negative | 0.56 | 0.39–0.80 | **0.001** | 0.68 | 0.45–1.03 | 0.071 | 0.65 | 0.42–1.00 | **0.047** |
| EBV positive | 0.61 | 0.12–3.08 | 0.548 | 1.28 | 0.15–11.3 | 0.823 | 0.49 | 0.10–2.44 | 0.382 |
| **p53 staining** | | | | | | | | | |
| Aberrant | 0.52 | 0.35–0.79 | **0.002** | 0.70 | 0.43–1.14 | 0.151 | 0.65 | 0.41–1.04 | 0.074 |
| Wild type | 0.75 | 0.37–1.53 | 0.430 | 0.73 | 0.34–1.56 | 0.421 | 0.77 | 0.33–1.80 | 0.550 |
| **ACRG** | | | | | | | | | |
| p53aber | 0.49 | 0.28–0.85 | **0.011** | 0.57 | 0.31–1.04 | 0.065 | 0.53 | 0.27–1.04 | 0.063 |
| p53wt | 1.04 | 0.37–2.86 | 0.948 | 0.82 | 0.28–2.40 | 0.713 | 1.23 | 0.035–4.37 | 0.752 |
| MSI | 0.77 | 0.36–1.67 | 0.508 | 0.95 | 0.38–2.37 | 0.911 | 0.43 | 0.15–1.24 | 0.119 |
| EMT | 0.51 | 0.25–1.01 | 0.053 | 0.61 | 0.24–1.60 | 0.317 | 0.92 | 0.46–1.85 | 0.815 |
| **TCGA** | | | | | | | | | |
| CIN | 0.34 | 0.13–0.86 | **0.023** | 1.06 | 0.35–3.19 | 0.922 | 1.25 | 0.48–3.23 | 0.644 |
| GS | 0.58 | 0.36–0.92 | **0.021** | 0.53 | 0.31–0.088 | **0.014** | 0.64 | 0.37–1.11 | 0.109 |
| MSI | 0.77 | 0.36–1.67 | 0.508 | 0.95 | 0.38–2.37 | 0.911 | 0.43 | 0.15–1.24 | 0.119 |
| EBV | 0.61 | 0.12–3.08 | 0.504 | 1.28 | 0.15–11.3 | 0.823 | 0.49 | 0.10–2.44 | 0.382 |

Abbreviations: CXCL12, C-X-C motif chemokine ligand 12; SCF, stem cell factor; HR, hazard ratio; CI, confidence interval; MMRp/d, mismatch repair proficient/deficient; EBVish, Epstein–Barr virus in situ hybridization; ACRG, Asian Cancer Research Group; p53aber/wt, p53 aberrant/wild-type; MSI, microsatellite instability; EMT, epithelial–mesenchymal transition; TCGA, The Cancer Genome Atlas; CIN, chromosomal instability; GS, genetically stable.
